# Common and dissociable contributions of alexithymia and autism to domain-specific interoceptive dysregulations – a dimensional neuroimaging approach

**DOI:** 10.1101/432971

**Authors:** Jialin Li, Lei Xu, Xiaoxiao Zheng, Meina Fu, Feng Zhou, Xiaolei Xu, Xiaole Ma, Keshuang Li, Keith M. Kendrick, Benjamin Becker

## Abstract

Alexithymia represents a transdiagnostic marker across psychiatric entities associated with emotional impairments, including autism spectrum disorders (ASD). Accumulating evidence suggests that interoceptive dysfunctions that underpin the core symptomatic emotion recognition and empathy deficits in ASD may be contributed to by high levels of alexithymia rather than autistic symptoms per se. However, previous findings are hampered by generally elevated alexithymia in ASD patients, and thus were not able to differentiate common and distinct contributions across the entire spectrum of variations of autism and alexithymia. Moreover, the multi-factorial nature of the domains affected, such as distinct neural reactivity towards perceiving physical and affective pain, has not been accounted for. Against this background the present fMRI study employed a dimensional trait approach in n = 242 healthy subjects to determine common and distinct associations between both traits and pain empathic responses towards physical and affective pain. Higher levels of alexithymia associated with increased left anterior insula pain empathic reactivity. Disentangling these effects revealed a positive association during perceived physical pain, but a negative one during affective pain. No significant associations with trait autism were found, but an interaction effect between the trait dimensions was observed in the mid-cingulate cortex. Moderation analysis demonstrated that trait autism only impacted mid-cingulate reactivity towards physical pain in high alexithymia subjects, whereas reactivity towards affective pain was specifically associated with trait autism in low alexithymia subjects. Findings confirm previous patient studies suggesting that alexithymia rather than autism per se may drive altered insula pain empathic reactivity. Importantly, the present approach allowed for the first time to demonstrate that the impact of alexithymia on insula reactivity varies as a function of the pain empathic domain and that effects on other core empathy nodes evolve in interaction with trait autism.

## Introduction

Alexithymia is characterized by impairments in identifying and describing ones’ internal feelings (Sifneos, 1973). Increased levels of alexithymia represent a transdiagnostic marker across psychiatric and psychosomatic entities associated with severe functional impairments in the emotional domain, with autism spectrum disorders (ASD) being a prime example (Terock et al., 2017; Valdespino et al., 2017). Whereas overarching frameworks propose that early disturbances in interoceptive processes may underpin the core symptomatic emotion recognition and empathy deficits in ASD (Quattrocki & Friston, 2014), accumulating evidence suggests that these impairments may be contributed to by high levels of alexithymia rather than autistic symptoms per se (see e.g. overview in Brewer et al., 2015). However, these initial findings in patient populations are hampered by generally elevated levels of alexithymia in ASD, and thus were not able to differentiate the common and distinct contributions across the entire spectrum of variations of autism and alexithymia. Moreover, the multi-factorial nature of interoception-associated functional domains, such as distinct neural pain empathic responses towards perceiving physical and affective pain in others (Lamm et al., 2011), has not been accounted for. Against this background the present fMRI study employed a dimensional trait approach in a large sample of n = 242 healthy subjects (122 males; mean age, 21.60 ± 2.35 years) to determine common and distinct associations between individual variations in both traits and the domain-specific pain empathic neural responses towards physical and affective pain.

## Methods and Materials

### Subjects

252 healthy, right-handed Chinese students participated in the experiment after providing written informed consent. Four subjects were excluded due to excessive head motion during fMRI acquisition (>3 mm or 3 degrees). Data from 6 subjects were lost due to technical failure. Consequently, data from n = 242 subjects was included in the final analyses (122 males; mean age, 21.60 ± 2.35 years). All subjects were free from current or a history of physical, neurological, or psychiatric disorders. Subjects were excluded in case they reported regular or current use of nicotine, alcohol, illicit drugs and medication. The study procedures were approved by the local ethics committee at the University of Electronic Science and Technology of China (UESTC) and in accordance with the revision of the Declaration of Helsinki.

### fMRI paradigm and stimuli

Pain empathy networks were assessed using a modified version of validated pain empathy paradigm. The block-design fMRI paradigm incorporated previously validated visual stimuli including pictures displaying painful everyday scenes from a first-person perspective (physical pain) and painful facial expressions (affective pain) as well as corresponding non-painful control stimuli for both stimuli types (Sheng & Han, 2012; Meng et al., 2012; Yao et al., 2016). All physical pain stimuli showed a person’s hand or foot in painful everyday situations from a first-person perspective (e.g. cutting a hand with a knife, the matched non-painful control stimulus shows cutting vegetables with a knife) (for stimuli see Meng et al., 2012). The affective stimuli consisted of painful and neutral expressions from 16 Chinese subjects (8 males) (for stimuli see Sheng & Han, 2012). In previous studies, both painful stimuli sets have been rated as more painful and increased activity in the pain empathy networks relative to the control stimuli (Sheng & Han, 2012; Meng et al., 2012; Yao et al., 2016).

For the present study, a total of 64 pictures were included (16 per experimental condition: physical pain/affective pain/physical control/affective control). The block design incorporated 16 blocks (4 blocks per condition) with 4 stimuli per block from the same category (each presented for 3s), and the blocks were interspersed by a jittered inter-block interval of 10s (8–12s). Total duration of the paradigm was 436s acquired in a single fMRI run. In order to minimize interference by cognitive processing, subjects were instructed to passively view the stimuli during scanning.

### Alexithymia and autism scales and quality assessment

To determine common and distinct contributions of alexithymia and autism to the neural pain empathic responses, levels of trait autism and alexithymia were assessed using validated Chinese versions of the Autism Spectrum Quotient (ASQ) (Baron-Cohen et al., 2001) and Toronto Alexithymia Score (TAS) (Bagby et al., 1994). Cronbach’s α scores in the present sample were 0.744 for ASQ and 0.817 for TAS. Assessing the normal distribution of the scales revealed that ASQ scores in the present sample were non-normal distributed (Shapiro-Wilk test, *p* < 0.05), to this end associations with neural indices were determined using non-parametric approaches (for a concordant approach see also Luo et al., 2018). ASQ and TAS scores were positively associated in the present sample (*r*ho = 0.408, *p* < 0.001), therefore the variance inflation factor (VIF) was assessed to test for problematic collinearity (Luo et al., 2018; Mumford et al., 2015). VIF in present study was 1.21, arguing against problematic collinearity (Mumford et al., 2015; see also Luo et al., 2018; Ohashi et al., 2017; Chau et al., 2017).

### Image Acquisition

MRI data were collected using a 3.0 Tesla GE MR750 system (General Electric Medical System, Milwaukee, WI, USA). A total of 218 functional volumes of T2*-weighted echo planar images were obtained for each subject using the following acquisition parameters: repetition time, 2000ms; echo time, 30 ms; slices, 39; slice-thickness, 3.4mm; gap, 0.6mm; field of view, 240 × 240 mm^2^; matrix size, 64 × 64; flip angle, 90°. High-resolution whole-brain volume T1-weighted images were additionally acquired to improve normalization of the functional images (spoiled gradient echo pulse sequence with oblique acquisition, acquisition parameters: repetition time, 6 ms; echo time, minimum; flip angle, 9°; field of view = 256 × 256 mm; acquisition matrix, 256 × 256; thickness, 1 mm; 156 slices). OptoActive MRI headphones (http://www.optoacoustics.com/) were used to reduce acoustic noise exposure for the participants MRI acquisition.

### MRI data analysis – preprocessing and analysis approach

fMRI data were analyzed using SPM12 software (Wellcome Trust Center of Neuroimaging, University College London, London, United Kingdom). The first ten volumes were discarded to (1) achieve magnet-steady images and (2) allow active noise cancelling by the headphones. The remaining functional images were realigned to correct for head motion, co-registered with the T1-weighted structural images and normalized using the segmentation parameters from the structural images to Montreal Neurological Institute (MNI) standard space. Normalized images were written out at 3mm^3^ voxel size and spatially smoothed using a Gaussian kernel with full-width at half-maximum (FWHM) of 8mm.

On the first level, the four experimental conditions, ‘physical pain’, ‘affective pain’, ‘physical control’ and ‘affective control’ were modeled using a box-car function subsequently convolved with the standard hemodynamic response function (HRF). The six head-motion parameters were included in the design matrix to further control for movement-related artifacts. Specific contrast images between painful and non-painful conditions were created for each subject (physical pain>physical control, affective pain>affective control). To examine associations with distinct brain networks engaged in physical and affective pain (Bruneau et al., 2012; Kanske et al., 2015; Richardson et al., 2018), the interaction contrast [(physical pain>physical control) > (affective pain>affective control)] was considered as main contrast of interest.

Given that the ASQ scores in the present sample were non-normal distributed (Shapiro-Wilk test, *p*<0.05), non-parametric tests were employed using the Statistical nonParametric Mapping toolbox (SnPM13, http://warwick.ac.uk/snpm) based on 10,000 random permutations to implement separate regression analysis for ASQ, TAS and their interaction (similar approach see also Luo et al., 2018). For the three regression models, one of the values was entered as predictor whereas the other two terms were entered as additional regressors of no interest. Results were thresholded at *p*<0.05 FWE-corrected on the cluster level. In line with recent recommendations for the control of false-positives in cluster-based correction methods (e.g. Eklund et al., 2016; Slotnick, 2017), an initial cluster-forming threshold of *p*< 0.001 (uncorrected) was employed. Analyses were restricted to voxels with a high probability to represent gray matter (SPM gray.nii> 0.3).

### Further analysis: extraction of parameter estimates and moderation

To further disentangle and visualize associations between the predictors and empathy related neural activity, parameter estimates were extracted from significant clusters using MarsBaR. The differences of associations between (physical pain>physical control) contrast and (affective pain>affective control) contrast were computed using percentile bootstrap (Wilcox, 2016) based on 10,000 bootstrap samples, which is used to compare dependent correlation coefficients. To further evaluate whether the interaction between alexithymia × autism predicts brain pain empathic responses, a moderation analysis was conducted using the SPSS 22 PROCESS macro (Preacher & Hayes, 2004).

## Results

Results demonstrated that higher alexithymia is associated with increased responses in the left anterior insula during pain empathy (peak MNI coordinates: [-36 –6 18], *k* = 109, *t*_238_ = 4.76, *p*_FWE-cluster_ = 0.045) (**Figure 1A**). To control for potential effects of age and gender, these variables were included as nuisance regressors. Results indicated that associations with activity in left insula remained robust (peak MNI coordinates: [-36 –6 18], *k* = 135, *t*_236_ = 4.62, *p*_FWE-cluster_ = 0.032).

Disentangling these effects by extraction of parameter estimates revealed a positive association with alexithymia scores during perceived physical pain, but a negative one during affective pain (**Figure 1B**; *p*s < 0.025; correlation difference *p* = 0.007, percentile bootstrap).

**Figure 1 – legend:**
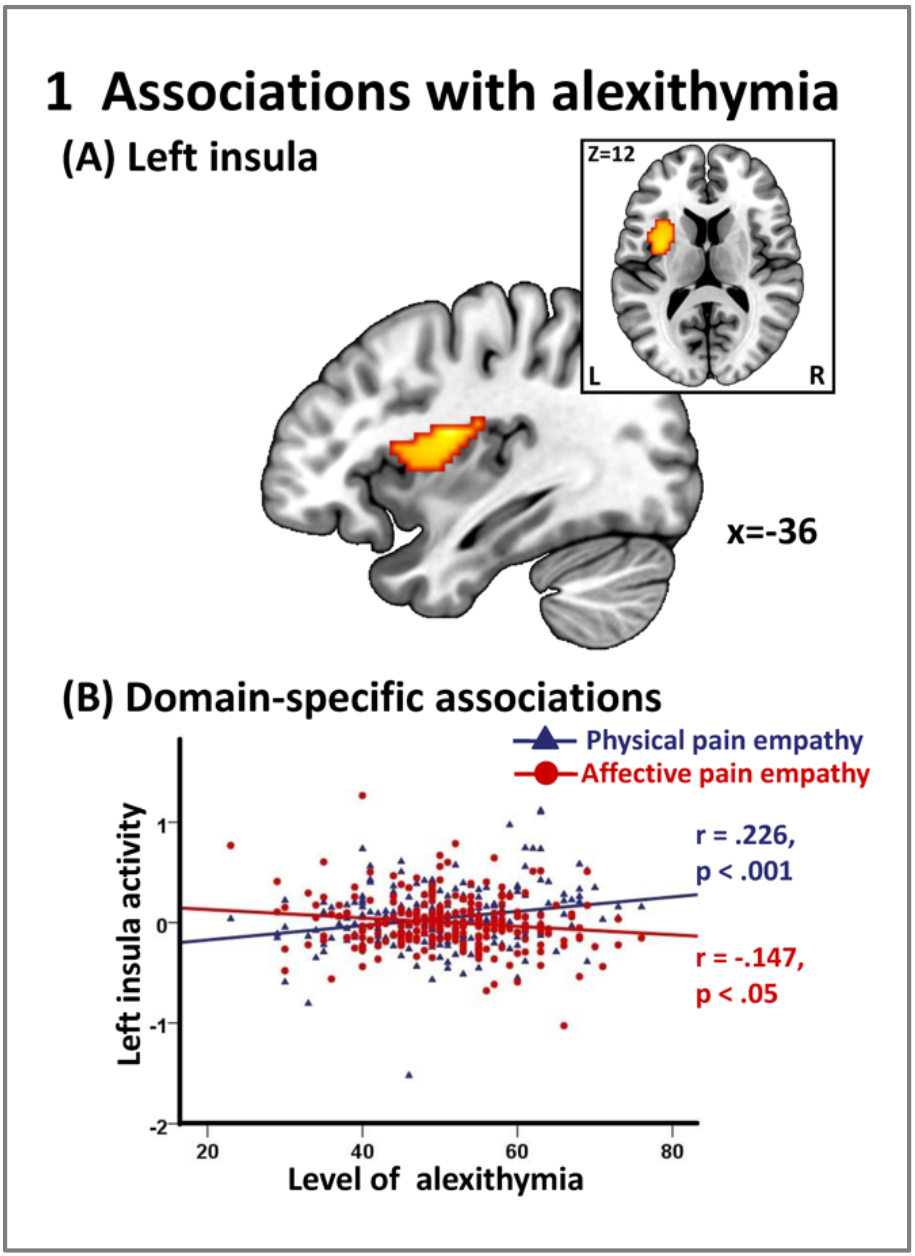
(**1A**) Levels of alexithymia were significantly associated with pain empathic responses in the left insula (peak MNI coordinates [-36, –6, 18], *k* = 109, *t*_238_ = 4.76, *p*_FWE-cluster_ = 0.045); (**1B**) extraction of parameter estimates from the left insula revealed that alexithymia showed opposite associations with insula reactivity towards physical and affective pain stimuli.

Whereas no significant associations with trait autism scores were found per se, there was an interaction effect between the two trait dimensions in the mid-cingulate cortex (peak MNI coordinates: [6 –6 36], *k* = 347, *t*_238_ = 4.46, *p*_FWE-cluster_ = 0.011) (**Figure 2A**). Associations remained stable after additionally controlling for gender and age as nuisance variables (peak MNI coordinates: [6 –6 36], *k* = 319, *t*_236_ = 4.41, *p*_FWE-cluster_ = 0.009). Extraction of parameter estimates demonstrated that the interaction term was positively associated with mid-cingulate reactivity during perceived physical pain, yet negatively with that for affective pain (**Figure 2B**; *p*s < 0.013; correlation difference *p* = 0.002).

**Figure 2 – legend:**
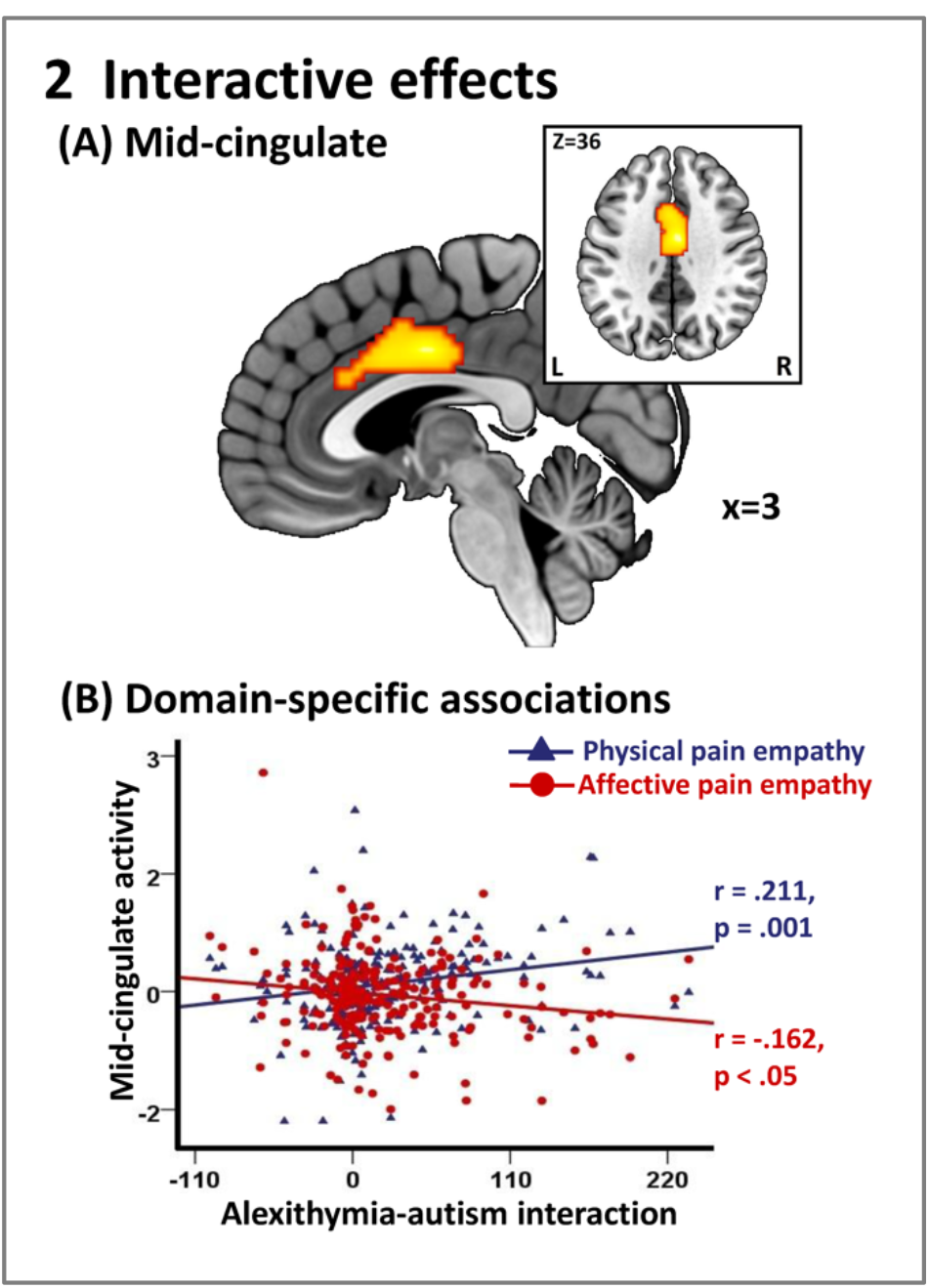
(**2A**) Interaction effect between alexithymia and autism on pain empathic response in the midcingulate cortex (peak MNI coordinates [ 6, –6, 36], *k* = 347, *t*_238_ = 4.46, *p*_FWE-cluster_ = 0.011); (**2B**) extraction of parameter estimates from the mid-cingulate revealed opposite associations during processing of physical versus affective pain.

To further disentangle the interaction effect we tested whether alexithymia moderated the impact of autism on mid-cingulate reactivity. Moderation analysis revealed significant moderation effects of alexithymia for mid-cingulate responses to both perceived physical and affective pain (physical, R^2^ = 0.058, *F*(3, 238) = 4.88, *p* = 0.003; affective, R^2^ = 0.048, *F*(3, 238) = 3.97, *p* = 0.009), revealing that its activation reflects an interaction between the traits (physical, *B* = 0.002, SE = 0.001, *t*_238_ = 3.28, *p* = 0.001; affective, *B* = –0.002, SE = 0.001, *t*_238_ = –2.59, *p* = 0.010). To further disentangle the effects, the moderator variable was split into three levels using the Johnson-Neyman approach, demonstrating that levels of trait autism only impacted mid-cingulate reactivity towards perceived physical pain in subjects with high alexithymia (**Figure 3A**; *t*_238_ = 2.80, *p* = 0.006), whereas mid-cingulate reactivity towards perceived affective pain was specifically associated with trait autism in subjects with low alexithymia (**Figure 3B**; *t*_238_ = 2.48, *p* = 0.014).

**Figure 3 – legend:**
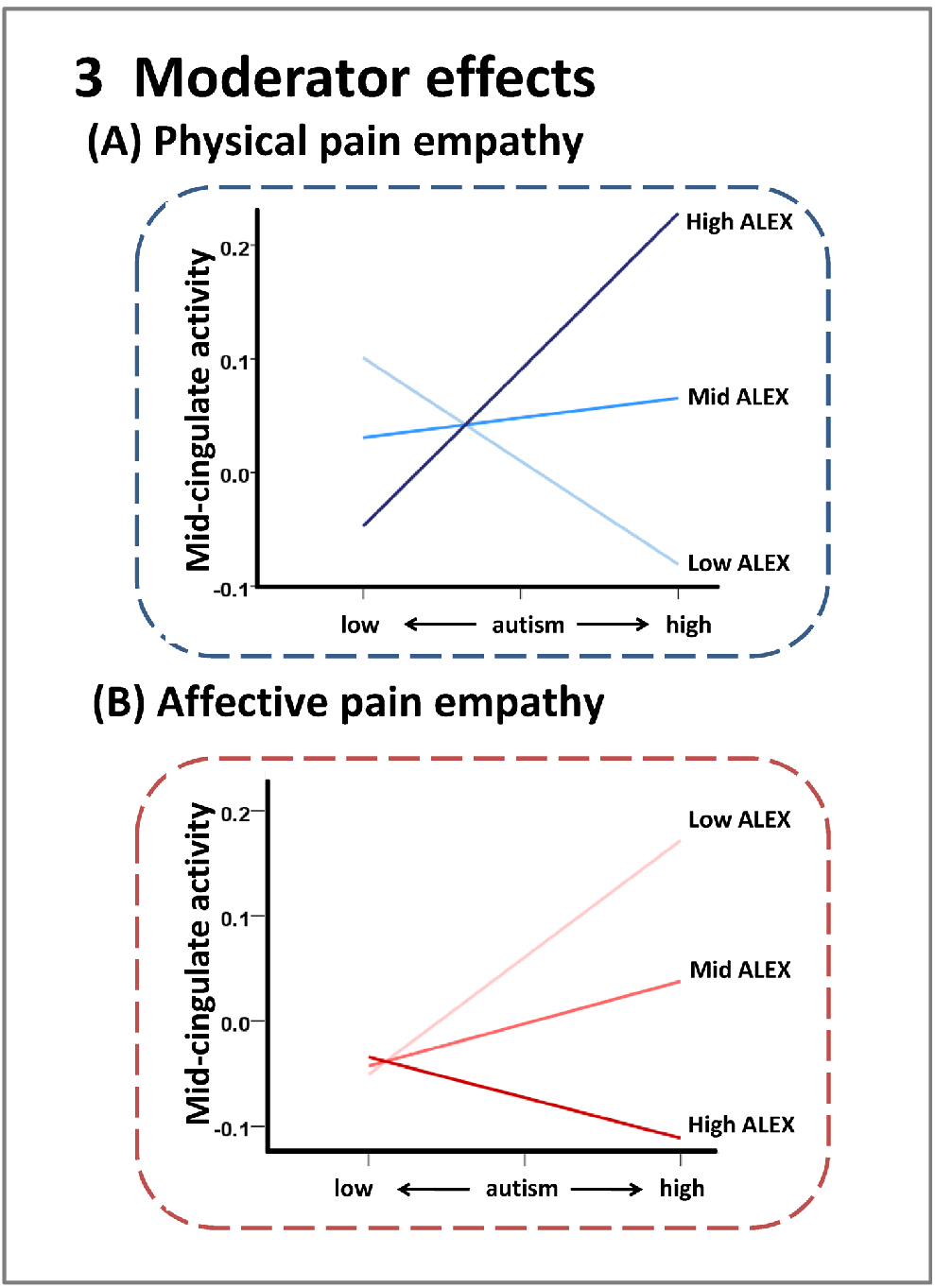
**(3A)** moderator effects of alexithymia (ALEX) on the association between autism and midcingulate reactivity towards physical and **(3B)** affective pain stimuli.

## Discussion

In line with research in ASD patients the present dimensional trait approach confirmed that alexithymia rather than autism per se drove altered insula interoceptive processing in the domain of pain empathy (e.g. Bird et al., 2010; for an overview see also Brewer et al., 2015; Valdespino et al., 2017). Importantly, employing a dimensional approach in combination with a multi-domain paradigm allowed to demonstrate for the first time that the impact of alexithymia on interoception-related insula processing varies as a function of physical versus affective pain empathy induction. The anterior insula is a functionally heterogenous region and interoceptive signaling within it not only underlies empathic responses but also inner affective experience and salience attribution to environmental stimuli (Uddin et al., 2017) including emotional faces (Yao et al., 2018). The present results indicate that higher levels of alexithymia may be associated with deficient attenuation of interoceptive signaling in response to physical pain signals, accompanied by under-responsivity towards socially transmitted pain signals. Given that early dysregulations in the oxytocin system may contribute to interoception-related deficits in ASD (Quattrocki & Friston, 2014) and oxytocin-administration can decrease insula responsivity to pain empathy (Bos et al., 2015), but increase it to social salience (Yao et al., 2018), the present findings may suggest that alexithymia is the moderating factor for both ASD-related deficits and oxytocinergic modulation of interoception-related insula processes.

Finally, employing a dimensional trait approach enabled to determine interaction effects of the two pathology relevant traits on mid cingulate empathic reactivity. The observed moderating role of alexithymia on the impact of autistic traits on mid-cingulate empathic reactivity supports a multi-factorial view on alixithymia’s contribution to ASD symptomatology. Together with the anterior insula, the mid-cingulate represents a key node for empathic processing as well as awareness of negative affect (Lamm et al., 2011; Tolomeo et al., 2016). The observed interaction suggests that alterations in the sub-domains further vary as a function of alexithymia with high values in both traits leading to exaggerated reactivity towards perceived physical pain, whereas aberrant responses to affective pain may represent a unique feature of alexithymia. Given the high symptomatic heterogeneity of ASD, concomitant assessments of alexithymia and sub-domain specific pain empathic responses may thus help to determine diagnostic- and treatment-relevant ASD subtypes.

## Financial disclosures and acknowledgements

This work was supported by the National Natural Science Foundation of China (NSFC, 91632117 to BB; 31530032 to KK), Fundamental Research Funds for the Central Universities of China (ZYGX2015Z002 to BB) and the Sichuan Science and Technology Department (2018JY0001 to BB). The authors report no biomedical financial interests or potential conflicts of interest.

